# Clinical Trial and Ontology-Derived Positive and Negative Benchmark Datasets for Drug Repurposing Across Rare Diseases

**DOI:** 10.64898/2026.06.15.732135

**Authors:** Cyrus Babak Ravandi, William R. Mowrey, Ayan Chatterjee, Fariba Khanshan, Parham Haddadi, Juan Carlos Mobarec, Simon Lambden, Tina Eliassi-Rad, Piero Ricchiuto, Gabriel Risa

## Abstract

Evaluating the potential applications of a medicine is a fundamental challenge in drug development. There is a lack of standardized, decision-oriented benchmarks that test whether computational models can generalize therapeutic hypotheses across diseases in ways that reflect real-world pharmaceutical investment decision making. To address this gap, we introduce two complementary resources: the Indication Expansion Investment Decision Network (IxIDN) and the Orphanet Rare Disease Ontology Negative-network (ORDON). IxIDN is a clinical-trial-derived positive benchmark constructed by projecting drug–disease associations from pharmaceutical clinical trials into a disease–disease network; each edge connects disease pairs that have entered clinical trials for the same drug, thereby capturing cases when concrete indication-expansion decisions have been made. The current release contains 574 rare diseases and 5,336 edges. In contrast, ORDON serves as a stringent, biology-aware negative benchmark derived from the authoritative Orphanet Rare Disease Ontology. It identifies maximally distant disease pairs according to curated hierarchical structure and genetics-linked inheritance patterns, providing 793 rare diseases and 5,000 edges that represent high-separation negative candidates across therapeutic areas. Together, IxIDN and ORDON enable rigorous cross-evidence generalization from clinical trials to disease ontology, testing for Disease– Disease Association Learning (DDAL), a core task for mechanism-centered drug repurposing and indication expansion. All data are publicly available with detailed metadata, enabling reproducible evaluation of models on transparent, decision-relevant benchmarks.

## Background & Summary

A fundamental challenge in translational medicine lies in the computational modeling of complex human diseases. At present, limited tools are available to support cross-disease generalization for indication selection or expansion. This adds risk and delay the development of new therapies for patients [1, 2]. Rare diseases, in particular, operate in a data-scarce environment that demands models capable of transferring mechanistic insights from better-studied non-rare disorders while also allowing unique biological findings from rare diseases to inform non-rare disorders [3–7].

Existing computational approaches to disease modeling offer complementary strengths but also face important evaluation challenges when applied to therapeutic decision-making. Knowledge graph and graph ML models [8–10] are prone to shortcut learning: by training and evaluating on the same curated graph, they inflate performance metrics while capturing label co-occurrence rather than shared biological mechanism — a problem serious enough that recent position papers have questioned the translational relevance of the entire graph learning paradigm [11, 12]. Large language models [13, 14] face a different failure: their disease associations are fluent but ungrounded. Because they encode statistical co-occurrence from text rather than causal pathway structure, they readily propose repurposing hypotheses that are narratively convincing yet mechanistically inert a drug-disease pair may look plausible in a sentence and be biologically meaningless in a cell [15, 16]. Agentic AI systems compound this problem [17]: hallucinated pathways, i.e., chains of reasoning that traverse plausible-sounding molecular steps with no physical substrate, propagate silently across retrieval and multi-step inference, making them particularly dangerous in a therapeutic context where a confident but false mechanistic argument can misdirect investment decisions [18–20].

These reframing shifts drug repurposing from label transfer to mechanism transfer and establish *Disease-Disease Association Learning (DDAL)* as a core task for translational AI. To address these gaps, we propose redefining disease-disease relatedness through *actionable therapeutic transferability*: two diseases are meaningfully related if a therapeutic hypothesis can plausibly transfer between them.

### Three axioms of mechanism-centered disease modeling

1. Diseases are not atomic classes; they are distributions of dysregulated biological pathways and processes shaped by context [21, 22]. The core hypothesis is that diseases sharing biological pathways and some phenotypic features are likely to converge on the same molecular targets even if their clinical manifestations differ. A canonical example is Crohn’s disease (CD) and ulcerative colitis (UC), two forms of inflammatory bowel disease (IBD) [23] that exhibit distinct pathology yet share non-rare symptoms and overlapping disease–gene associations.
2. Repurposing is mechanism transfer, not label transfer [24, 25]; In other words, label transfer means simply assigning a drug to a new disease because both share a broad clinical label (e.g., both are described as “autoimmune disorders”), without requiring any shared underlying biological mechanisms. In contrast, mechanism transfer requires that the diseases share dysregulated pathways or processes that the drug can actually target. While most disease ontologies are primarily etiology- and label-driven, real clinical medicine defines diseases through patient phenotypes, biomarkers, and population-specific manifestations [26].
3. Therefore, the learning target must include disease-disease therapeutic transferability [27, 28], not only drug ↔ disease associations.

To address this need for decision-oriented, mechanism-centered benchmarks, and pave the way toward developing models that follow the aforementioned axioms, we propose two complementary benchmarks: (i) the Indication Expansion Investment Decision Network (IxIDN) as positive samples, and (ii) the Orphanet Rare Disease Ontology Negative-network (ORDON) as negative samples. In the following, we demonstrate how the IxIDN–ORDON benchmarks can be used to establish rigorous cross-evidence generalization tests for assessing large language models (LLMs), agentic AI systems, and heterogeneous knowledge-graph machine learning (GraphML) models on mechanism-based therapeutic transferability tasks.

## Methods

### Data Preparations

#### Indication Expansion Investment Decision Network (IxIDN) as Positive Samples

We introduce IxIDN constructed directly from pharmaceutical clinical trial data to serve as a positive benchmark for DDAL as a proxy for disease–disease therapeutic transferability. A bipartite disease–drug graph was derived from clinical trial records, where an edge represents at least one trial investigated a drug–disease pair. An unweighted projection was then applied to transform this bipartite graph into a disease–disease network. Each edge in IxIDN connects a pair of diseases for which the same drug has entered clinical trials for both indications, thereby representing at least one concrete investment decision and a hypothesis of shared biological mechanism suitable for indication expansion (Figure 1A). The current release of IxIDN contains 574 diseases and 5,336 edges. IxIDN captures the complexity of indication expansion investment decisions evaluated in clinical trials.

**Figure 1:**
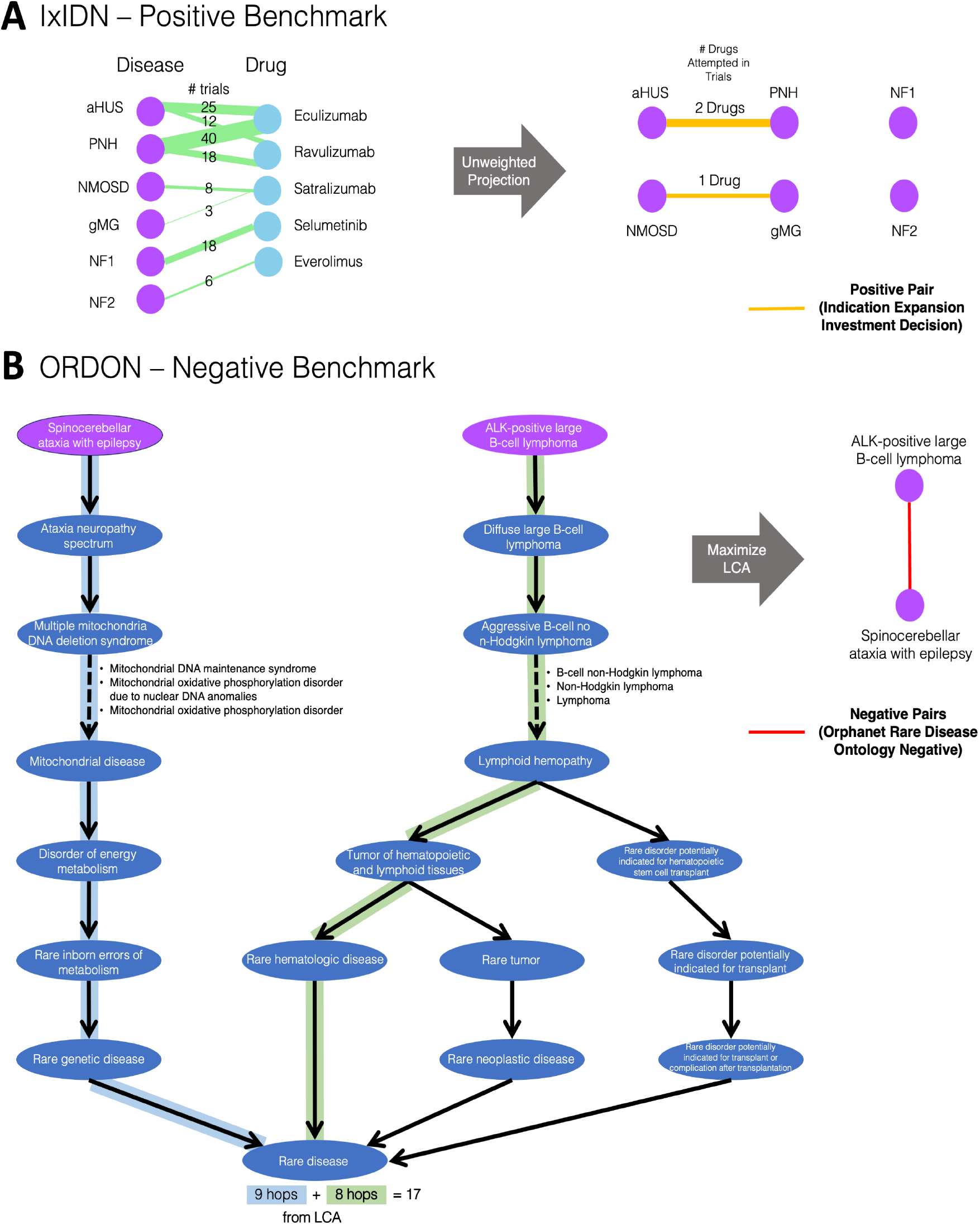
Overall evaluation paradigm for Disease-Disease Association Learning (DDAL) using the IxIDN and ORDON benchmarks. **(A) IxIDN – Benchmark for Positive Links**. This panel illustrates the *association-based framing of therapeutic transferability through the two-step mechanistic linkage*: the projection of drug ↔ mechanism evidence to mechanism ↔ disease-state association. The left side shows a bipartite disease–drug graph derived from clinical trial data, with edges indicating the number of trials per drug–disease pair. Through unweighted projection (gray arrow), this is transformed into a disease–disease network (right side). Yellow edges represent positive pairs corresponding to real-world indication expansion investment decisions (i.e., the same drug has entered clinical trials for both diseases). **(B) ORDON – Benchmark for Negative Links**. The left panel shows the construction of stringent negative samples by maximizing the LCA distance within the ORDO hierarchy. The right panel highlights an example negative pair (red edge) between Spinocerebellar ataxia with epilepsy and ALK-positive large B-cell lymphoma, separated by total of 17 hops from max LCA, this is one of the most distant pairs in the ORDO hierarchy. This biology-aware negative benchmark reflects clear mechanistic boundaries across therapeutic areas in rare diseases.

#### Orphanet Rare Disease Ontology Negative-network (ORDON) as Negative Samples

To provide stringent and biology-aware negative samples, we introduce ORDON derived from the authoritative Orphanet Rare Disease Ontology (ORDO). Disease pairs were selected by maximizing the Lowest Common Ancestor (LCA) distance within the curated hierarchical structure of ORDO, which incorporates genetics-linked inheritance patterns that delineate clear mechanistic boundaries across therapeutic areas. An example negative pair is Spinocerebellar ataxia with epilepsy and ALK-positive large B-cell lymphoma, separated by a total of 17 hops from the maximum LCA (Figure 1B). The current release of ORDON contains 793 diseases and 5,000 edges, representing a diverse pair of biologically distant rare diseases. Because ORDO spans multiple therapeutic areas, these maximally distant pairs naturally cross therapeutic-area boundaries and represent strong mechanistic separation, reflecting clear mechanistic boundaries across therapeutic areas, even though all pairs are selected from rare diseases.

To further strengthen the quality and biological stringency of the ORDON negative benchmark, we examined the full distribution of summed LCA distances across all disease pairs in the ORDO hierarchy. From this distribution, we selected top 5,000 pairs in the extreme right tail (Figure 2), corresponding to the most mechanistically distant pairs as stringent negative samples. This selection ensures maximal separation across therapeutic areas while remaining fully grounded in the curated ontology structure.

**Figure 2:**
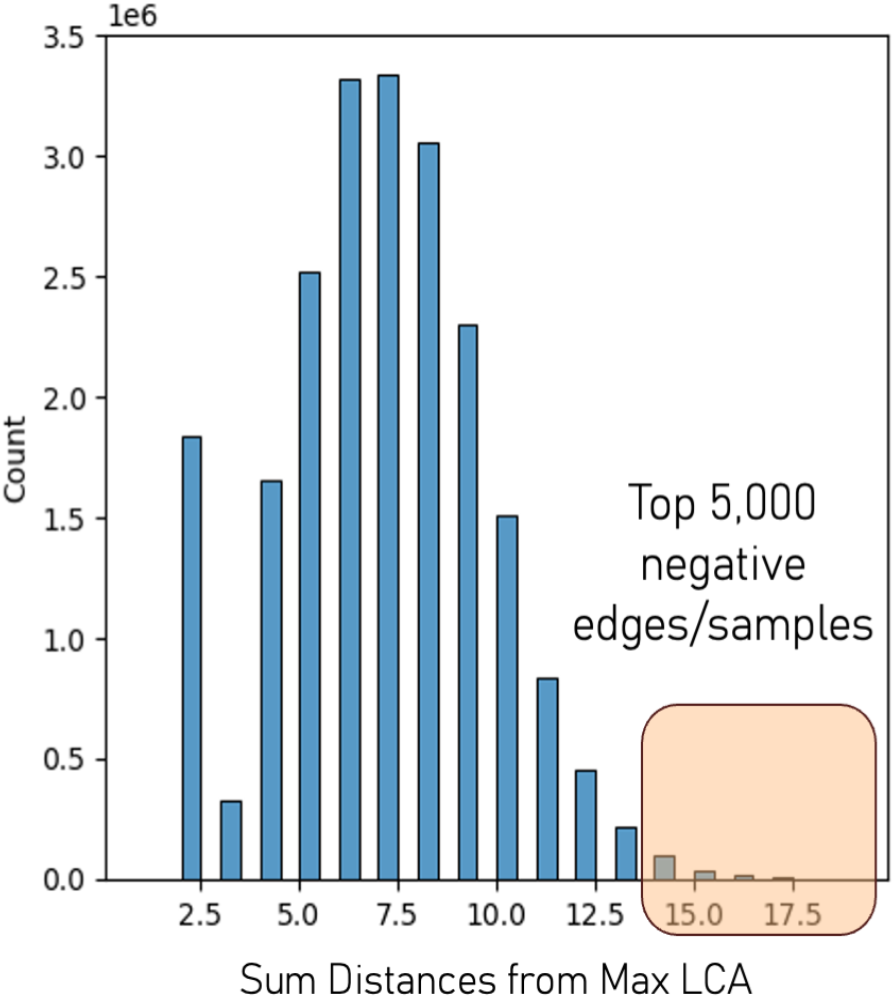
Distribution of summed Lowest Common Ancestor (LCA) distances for all disease pairs in the ORDO hierarchy. The histogram shows the frequency distribution of pairwise summed LCA distances across all disease pairs. To mitigate class imbalance between positive samples (IxIDN) and negative pairs, we selected the top 5,000 most distant pairs (highlighted in orange in the extreme right tail). These pairs were used as stringent negative samples to construct the ORDON benchmark.

### Data Record

All engineered datasets generated in this study are available through Zenodo: https://zenodo.org/records/20694608. The primary resource, IxIDN_ORDON_benchmark.csv, is a single CSV file containing the combined and blinded benchmark for reproducible evaluation. The accompanying files, IxIDN_df.csv and ORDON_df.csv, provide the source data used to construct the IxIDN and ORDON benchmarks. Figure 3 presents the degree distributions and summary statistics for IxIDN and ORDON. These distributions show that the two networks are structurally distinct, making them a useful benchmark for assessing the ability of DDAL models to distinguish between positive and negative predictions.

**Figure 3:**
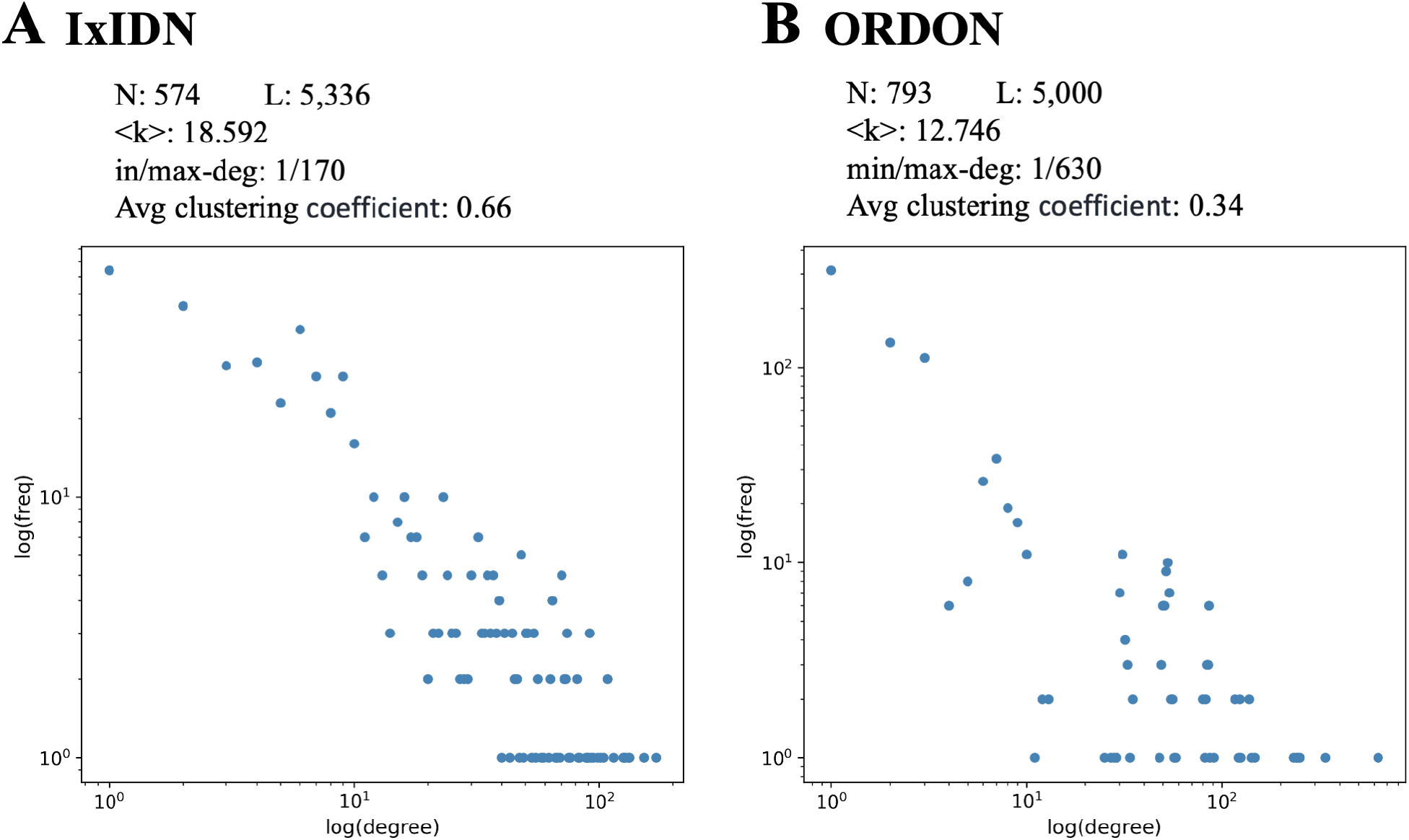
Network properties of the IxIDN and ORDON benchmarks. Scatter plots showing the degree distribution of (A) the IxIDN positive benchmark (574 nodes, 5,336 edges) and (B) the ORDON negative benchmark (793 nodes, 5,000 edges). Summary statistics are displayed above each panel (N = number of nodes, L = number of edges, ⟨k⟩ = average degree, min/max-deg = minimum and maximum node degree, and average clustering coefficient). Both networks resemble heavy-tailed properties with a broad range of node degrees.

- **IxIDN_ORDON_benchmark.csv**: The main benchmark file for disease–disease association learning evaluation. Columns: d1_disease_label (label of the first disease), d2_disease_label (label of the second disease). The file contains 10,336 randomly shuffled rows: 5,336 positive samples and 5,000 negative samples. This file enables direct use in validation of link-prediction and cross-evidence generalization tasks.
- **IxIDN_df.csv**: The file contains 574 rare diseases and 5,336 positive edges/samples, including rare disease IDs mapped to MONDO terminology, repurposed drug names, and the total number of trials aggregated to derive each investment decision link.
- **ORDON_df.csv**: This file contains 793 rare diseases and 5,000 negative edges/samples generated by maximizing LCA distances in ORDO, along with the Orphanet rare disease IDs associated with each negative link.

### Technical Validation

To validate the technical quality, biological relevance, and inherent difficulty of the IxIDN– ORDON benchmarks, we systematically evaluated the cross-evidence generalization capabilities of several representative, state-of-the-art artificial intelligence and GraphML models. By testing these models on the DDAL task using IxIDN and ORDON as validation datasets, we assess their capacity to recover therapeutic-transfer signals represented by the IxIDN and ORDON benchmarks. Collectively, these evaluations establish robust dataset-level sanity checks and provide reproducible performance baselines to anchor future comparative studies utilizing this benchmark framework. To rigorously evaluate these models, we implemented four distinct validation protocols:

- **Protocol 1: Large Language Model (LLM) Inference**. Evaluation of advanced language models configured in both parametric scoring modes (leveraging internal weights) and tool-augmented, agentic inference modes. In this protocol, we evaluated GPT-5.5 [29], Claude Opus 4.7 [30], and Gemini 3 [31].
- **Protocol 2: Iterative Agentic Retrieval and Reasoning**. Assessment of agentic AI systems that employ iterative multi-step retrieval-augmented generation (RAG) and reasoning over structured biomedical knowledge bases. In this protocol, we evaluated Edison Scientific [32, 33] and Phylo [34] platforms.
- **Protocol 3: Zero-Shot Knowledge Graph Foundation Model**. Evaluation of TxGNN [35], a foundation GraphML for clinician-centered drug repurposing, based on a knowledge graph. A modified TxGNN is assessed with drug nodes strategically withheld during training to prevent data leakage.
- **Protocol 4: Indirect Link Prediction via Target Overlap**. An indirect evaluation framework assessing disease–drug link prediction performance by quantifying the predicted drug-set overlap between the disease pairs defined in the IxIDN benchmark. In this protocol, we evaluated DeepDR (a deep autoencoder-based drug repositioning framework) [36], HeTDR (a heterogeneous graph transformer) [37], and HGTDR (a heterogeneous graph transformer with dual relational attention) [38].

To ensure unbiased evaluation, Protocols 1 and 2 were conducted under strict blinding of the ground-truth class labels. The IxIDN_ORDON_benchmark.csv file was randomly shuffled, with true labels and disease IDs removed prior to evaluation. These blinded disease pairs were then passed to the agentic systems via a standardized prompt, instructing the models to compute and output continuous Disease–Disease Association (DDA) scores. While completely purging historical clinical trial data and drug-related associations from the pre-training corpora of proprietary large language models presents an inherent challenge, we mitigated potential data leakage and confounding biases by explicitly constraining the models’ reasoning pathways. Specifically, the prompt contains strict programmatic directives asking the agent to prohibit the utilization of clinical trial outcomes or existing drug indication data during the inference process. The exact prompt executed for these evaluations is detailed below:

> *“You are a disease-disease association learning (DDA) agent. Your task is to calculate DDA(d1, d2), which returns an association score between 0 and 1 for two diseases to quantify the extent these diseases share mechanisms of action. Strictly, do not use clinical trials and drug datasets to implement the DDA function*.
>
> *For each disease pair in the attached csv file, compute DDA and add a new column to the csv file named DDA to export the DDA scores for each row. Make sure in the final csv file you are producing exact same columns as attached without any modification and just add a single new column named DDA.”*

Protocol 3 evaluates Graph Machine Learning (GraphML) approaches, framing DDAL as a link prediction task between disease pairs. In contrast to Large Language Models (LLMs), minimizing data leakage within GraphML architectures can be achieved with greater experimental control, as clinical trials and drug-related edges can be systematically pruned from the underlying training graph. To leverage this advantage, we evaluated TxGNN, a foundation model optimized for clinician-centered drug repurposing, by completely withholding drug nodes from its knowledge graph during a full retraining cycle.

Figure 4 reports the Area Under the Receiver Operating Characteristic curves (AUROC) and the empirical distributions of the computed continuous DDA scores across Protocols 1–3, quantifying the performance separation between the positive and negative classes. Strikingly, when data leakage is strictly controlled during training, TxGNN performance degrades to near-random levels. These results demonstrate that while mitigating leakage in LLMs remains an ongoing challenge, the IxIDN and ORDON benchmarks successfully differentiate the capacities of these diverse model classes to predict real-world drug repurposing investment decisions.

**Figure 4:**
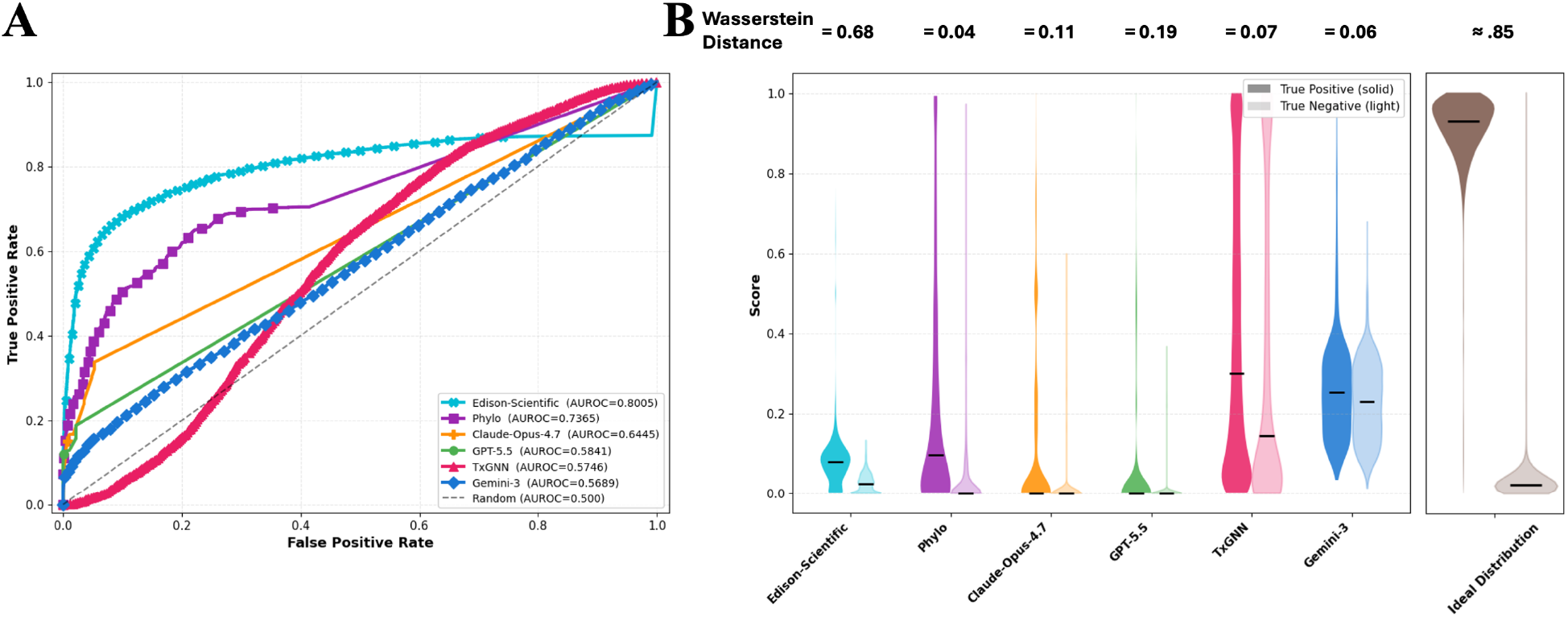
Reference baseline performance on the IxIDN–ORDON benchmark. (A) Receiver-operating-characteristic (ROC) curves and (B) The probability distributions of the normalized DDAL prediction scores between [0,1] for representative methods (large language models, agentic AI systems, and disease–drug knowledge-graph models) evaluated under the cross-evidence generalization protocols. These plots serve as dataset-level sanity checks, demonstrating clear separation between positive (IxIDN / NN) and negative (ORDON / NP) pairs while confirming that the benchmark presents a realistic and non-trivial challenge for current translational AI approaches.

Beyond establishing a robust benchmark for training and validating machine learning (ML) models for the DDAL task, we systematically analyzed differences in prediction distributions between the IxIDN and ORDON subsets. Specifically, we used the Wasserstein distance [39] to quantify the statistical divergence between the continuous prediction distributions of positive and negative samples. As shown in Figure 3, IxIDN and ORDON exhibit distinct degree distributions, indicating that the two networks differ substantially in their structural properties. In this setting, an ideal DDAL model would assign predictions close to 1 for positive links and close to 0 for negative links, producing strong separation between the two classes. Clear separation would correspond to maximal discriminative capacity, as illustrated in Figure 4B. Evaluating models under this expectation creates a rigorous benchmark challenge for both agentic and GraphML architectures, providing a useful stress test for reliability in translationally relevant biomedical applications.

The baseline analyses demonstrate clear statistical separation between IxIDN positives and ORDON negative samples, confirming that the benchmark captures biologically and clinically meaningful therapeutic-transfer signals. At the same time, performance remains moderate (well below perfect separation), indicating that the dataset presents a realistic and non-trivial challenge for current translational AI approaches. These reference results therefore validate the utility of the released resource for rigorous evaluation of mechanistic generalization in drug repurposing and indication-expansion research.

As an additional reference baseline, Protocol 4 evaluates three representative disease– drug knowledge graph models on the IxIDN benchmark: DeepDR [36], HeTDR [37], and HGTDR [38]. While these architectures were originally developed for drug–disease ranking tasks on heterogeneous knowledge graphs, we implemented an indirect evaluation framework to test whether they implicitly capture the therapeutic transferability signals encoded in real-world clinical trial investment decisions. Specifically, for each positive IxIDN edge connecting diseases *d*_1_ and *d*_2,_ we retrieved the top-10 predicted drugs ranked by each model for both diseases and quantified the size of their intersection.

The resulting distribution of intersecting-drug counts across all IxIDN edges is illustrated in Figure 5. Despite the diversity of the evaluated GraphML architectures and their exposure to historical clinical-trial disease-drug pairs, performance on this indirect task suggests limited recovery of the therapeutic-transfer signals represented in IxIDN. These results are consistent with the possibility that some disease-drug KG models rely substantially on explicit training topology, underscoring the value of complementary benchmarks that test generalization beyond observed graph structure.

**Figure 5:**
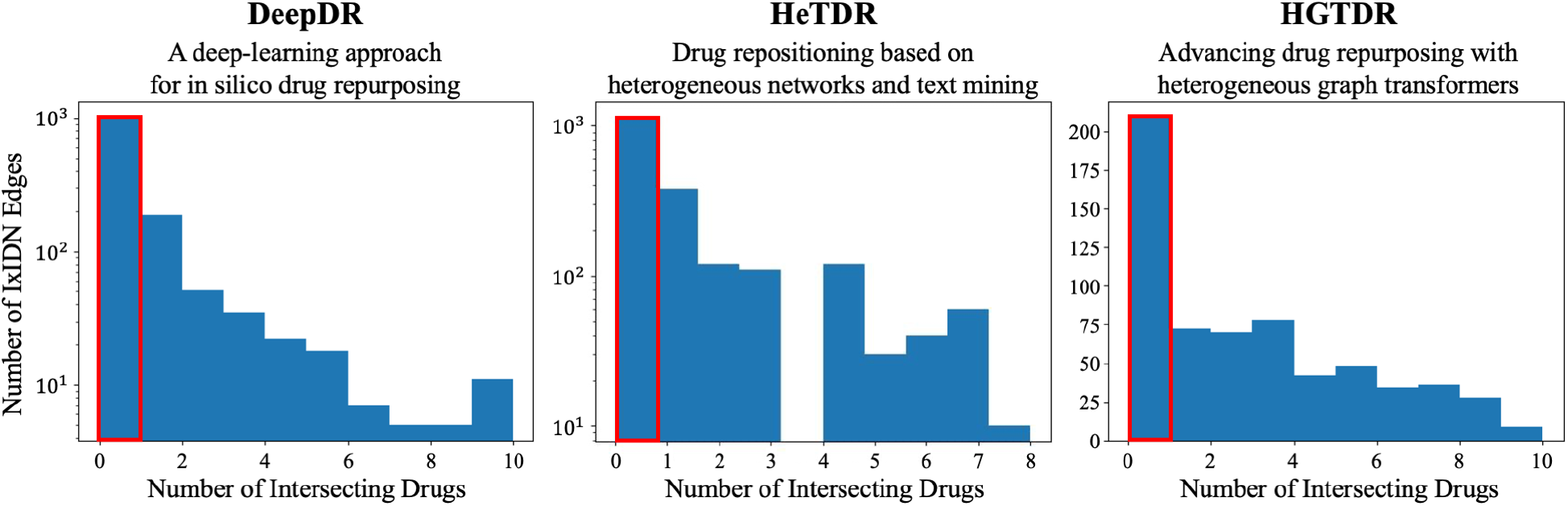
Indirect DDAL evaluation via drug-set overlap for selected disease–drug knowledge-graph models. Bar plots showing the distribution of the number of intersecting (shared) drugs between disease pairs in the IxIDN–ORDON benchmark for (A) DeepDR, (B) HeTDR, and (C) HGTDR. These plots provide an additional reference baseline that further illustrates the utility of the released dataset for evaluating therapeutic transferability through indirect drug-overlap approaches.

### Usage Notes

The IxIDN-ORDON benchmark is intended for evaluating computational models that score disease-disease therapeutic transferability, including link-prediction models, biomedical knowledge graph models, representation learning methods, and language-model-based biomedical reasoning systems. The combined benchmark file provides disease-pair labels suitable for binary classification or ranking tasks. IxIDN positive pairs should be interpreted as clinical-trial-derived evidence of prior therapeutic co-investigation, not definitive proof of shared mechanism. ORDON negative pairs should be interpreted as stringent ontology-derived negative candidates with high hierarchical separation, not absolute evidence that no biological or therapeutic relationship exists.

Users should avoid training and testing on overlapping clinical-trial-derived drug-disease edges, as this may inflate performance through data leakage. For fair evaluation, models should be assessed under conditions that restrict direct access to the clinical-trial evidence used to construct IxIDN. Researchers may also use the individual IxIDN_df.csv and ORDON_df.csv files to inspect source-specific metadata, disease identifiers, and construction logic before applying the shuffled combined benchmark.

## Code Availability

Custom code used for dataset construction and validation is not publicly available at this time. The released benchmark files, including IxIDN_ORDON_benchmark.csv, IxIDN_df.csv, and ORDON_df.csv, are available through Zenodo: https://zenodo.org/records/20694608, enabling reuse of the benchmark for downstream evaluation.

## Data Availability

IxIDN and ORDON are available through Zenodo: https://zenodo.org/records/20694608. These benchmarks were derived from Orphanet and Open Targets, both of which are publicly available resources that are regularly curated and updated. Orphanet provides rare disease ontology and terminology, while Open Targets provides public annotations linking diseases, clinical trials, and drugs.

## Acknowledgements

The authors thank Marek Justyna of the Institute of Computing Science, Poznan University of Technology, for his valuable contributions and fruitful discussions on benchmark evaluation. The authors also acknowledge AstraZeneca’s Scientific Computing Platform (SCP) for providing the computational infrastructure and resources that enabled this research.

## Competing Interests

Alexion AstraZeneca Rare Disease is a pharmaceutical company focused on developing novel therapeutics for rare diseases. A.C. is the founder of BioClarity AI, Inc., a company utilizing graph machine learning for applications in gene therapy. C.B.R, W.R.M., F.K., P.H., J.C.M., S.L., P.R., and G.R. are employees of AstraZeneca and may hold stock options.

Author Contributions

B.R., W.R.M., A.C., and P.R. conceived and designed the project. B.R., W.R.M., and A.C. wrote the manuscript and performed data curation and preparation. B.R. led data engineering and integration, designed the graph machine learning framework, developed the IxIDN and ORDON benchmark datasets, conducted all benchmarking experiments including evaluations on LLMs and foundation models, and performed network-based modeling and analysis. W.R.M. conducted an in-depth evaluation of the disease clusters and the investment decision benchmark. A.C. and B.R. designed the graph machine learning architecture and tuned the hyperparameters. F.K. provided clinical insights and contributed to manuscript writing. P.H. performed biological validations and contributed to manuscript writing. J.C.M. led the design of multiple case studies for validation of knowledge-graph approaches and made significant contributions to manuscript writing. S.L. provided clinical feedback and made substantial contributions to manuscript writing. T.E.R. provided guidance on machine learning experimental design and offered fundamental contributions to manuscript writing. P.R. contributed to overall study design and manuscript writing. G.R. contributed to the design and strengthening of the IxIDN and ORDON benchmarks, supported the maturation and execution of the Dis2Vec codebase, and played a key role in the validation, interpretation, evaluation of the benchmarks on LLMs and foundation models, and made significant contributions to manuscript writing.

